# Chronic starvation induces non-canonical pro-death stress granules

**DOI:** 10.1101/317529

**Authors:** Lucas C. Reineke, Shebna A. Cheema, Julien Dubrulle, Joel R. Neilson

**Affiliations:** Department of Molecular Physiology and Biophysics; Graduate Program in Integrative Molecular and Biomedical Sciences; Integrated Microscopy Core, Baylor College of Medicine. Houston, TX, USA

**Keywords:** Stress Granule, eIF2α, Ras-GAP SH3 binding protein, starvation, Translation, cell death

## Abstract

Stress granules (SGs) assemble under stress-induced conditions that inhibit protein synthesis, including eIF2α phosphorylation, inhibition of the RNA helicase eIF4a, or inactivation of mTORC1. Classically defined SGs are composed of translation initiation factors, 40S ribosomes, RNA binding proteins, and poly(A)^+^ mRNAs, and as such represent an important compartment for storage of mRNAs and regulation of their translation. Emerging work on SGs indicates they may play important roles in cancer, neurodegenerative disease, and viral infection, often promoting survival. Yet much previous work on SGs formation and function has employed acute stress conditions, which may not accurately reflect the chronic stresses that manifest in human disease. We used prolonged nutrient starvation to investigate SG formation and function during chronic stress. Surprisingly, SGs that form under chronic nutrient starvation lack 40S ribosomes, do not actively exchange their constituent components with cytoplasmic pools, and promote cell death. These results imply that SG assembly and function in the context of prolonged nutrient starvation stress differ significantly from what has been described for acute stress conditions.

**Summary Statement:** This work characterizes the mechanisms of formation of a novel type of stress granule that is induced in response to long-term starvation, and unlike previously described stress granules, functions in a pro-death capacity.

## Introduction

Stress granules (SGs) are RNA granules that contain translation initiation factors, RNA binding proteins, poly(A)^+^ RNA, and 40S ribosomes (Reineke & Lloyd 2013; Kedersha & Anderson 2007). Stress granules assemble due to accumulation of stalled translation initiation complexes that can be induced by phosphorylation of eukaryotic translation initiation factor 2α (eIF2α) or inhibition of the RNA helicase eIF4a (Kedersha et al. 2013; Dang et al. 2006; Kedersha et al. 2002). eIF2α phosphorylation is a well-established precursor to SG assembly, and is regulated by the antagonistic functions of eIF2α kinases and phosphatases. Four eIF2α kinases have been identified: HRI, PERK, PKR and GCN2. HRI is well characterized to be activated under conditions of low heme in erythrocytes, but also can act in response to oxidative stress such as is experienced during arsenite exposure (McEwen 2005). PERK primarily signals a state of endoplasmic reticulum (ER) stress (McQuiston & Diehl 2017). PKR is activated in response to cytoplasmic double-stranded RNA during viral infection and may respond to other physiological stressors including oxidative and ER stress, and cytokine signaling (Garcia et al. 2006). Finally, GCN2 is activated by amino acid deprivation and UV stress (Deng et al. 2002; Aulas et al. 2017).Thus, while each kinase is activated in response to a unique type of cellular stress, some commonality among their activation has been shown to exist based on the type and duration of the stress. Two phosphatase complexes have been shown to ameliorate the function of the eIF2α kinases, one containing GADD34 and the other containing CrEP (Harding et al. 2009; Hetz et al. 2013). The regulation of these phosphatase complexes is not well characterized.

Stress granule assembly has been studied extremely well in the context of acute stressors, including oxidative stress, ER stress, heat shock, and UV irradiation. Stress granules induced by acute stress contain, by definition, translation initiation factors, 40S ribosomes and RNA binding proteins such as nucleating proteins G3BP1 and G3BP2, Tia1 and TiaR. Some other stress granule-resident RNA binding proteins include HuR, PABP, and FMRP. Acute stress granules are also extremely dynamic, and recent work characterizing core complexes within acute SGs has led to a model of a dynamic shell, from which various components are exchanged with the cytosol, surrounding a stable core complex (Jain et al. 2016). Acute stress granules also contain many signaling molecules and recent evidence suggests that they might modulate signaling through several pathways including mTORC1, NFκB and innate immune signaling through PKR and RIG-I (Sfakianos et al. 2018; Reineke et al. 2015; Kim et al. 2005; Onomoto et al. 2012). Stress granule assembly during acute stressors has also been shown to regulate MAPK signaling resulting in the paradigm that stress granules function in a pro-survival response (Arimoto et al. 2008). This paradigm has resulted in the assumption that all stress granules function in this capacity despite findings indicating that SGs forming in the broad range of acute stressors vary in composition (Aulas et al. 2017).

Given the above pathways that promote SG condensation and the paradigm that SG function as pro-survival, we endeavored to investigate whether this paradigm also applies to chronic stress. We utilized nutrient starvation to model chronic stress because of its implications for numerous disease states including tumor biology, cardiac ischemia and stroke. Nutrient starvation has been shown to inhibit protein synthesis through mTORC1 inhibition, which has been implicated in acute stress granule formation. Therefore, studying nutrient starvation also provides insight into crosstalk of mTORC1 signaling and eIF2α phosphorylation in promoting SG assembly. We characterized SG dynamics and composition, and evaluated SG function in regulating cell fate during chronic nutrient starvation. Our findings indicate that novel stress granules form under conditions of chronic nutrient starvation stress. In contrast to canonical acute SGs, these novel RNA granules are less dynamic, do not contain 18S small ribosomal RNA, and act to promote cell death. Together these findings indicate that different SGSGs function differently and the paradigm of SG as pro-survival may not apply to all stress conditions.

## Results

We evaluated the formation of SGs in U2OS cells, utilizing a model of chronic nutrient starvation in which cells were starved of glucose, serum, glutamine and pyruvate. We first performed time-course experiments to investigate formation of SGs in response to these chronic nutrient starvation conditions. Strikingly, using the subcellular localization and aggregation of the SG nucleating RNA binding protein G3BP1 and eIF3b as canonical markers of SGs, SGs did not appear until 8h of nutrient starvation (Fig. 1A, B). These kinetics are far slower than the kinetics previously described for acute stressors (Kedersha et al. 2002; Kedersha et al. 2000; Farny et al. 2009; Aulas et al. 2017). Assessment of bulk translation using ribopuromycylation, as well as P-eIF2α and P-4EBP1 levels, revealed that while 4EBP1 phosphorylation at T37/46 and S65 decreases rapidly early in the stress, SG assembly itself closely correlated with the kinetics of eIF2α phosphorylation (Fig.1C). These results suggest that SG assembly under chronic nutrient starvation is not directly dependent on decreased mTORC1 activity but is likely to instead occur in response to eIF2α phosphorylation. Since acute SGs are known to depend on ATP for assembly and dynamics (Jain et al. 2016), and our starvation conditions were anticipated to deplete cells of ATP, we measured ATP levels using a luminescence-based assay. At 8h of starvation, concomitant with SG appearance, ATP levels had fallen to 25 % of that measured in fed controls (Fig.1D). After 16 h of starvation, the point at which all cells were consistently characterized as containing SGs, ATP levels were approximately 1 % of fed controls. This indicates that the observed SGs assemble coincident with declining ATP levels (Fig.1D).

**Figure 1:**
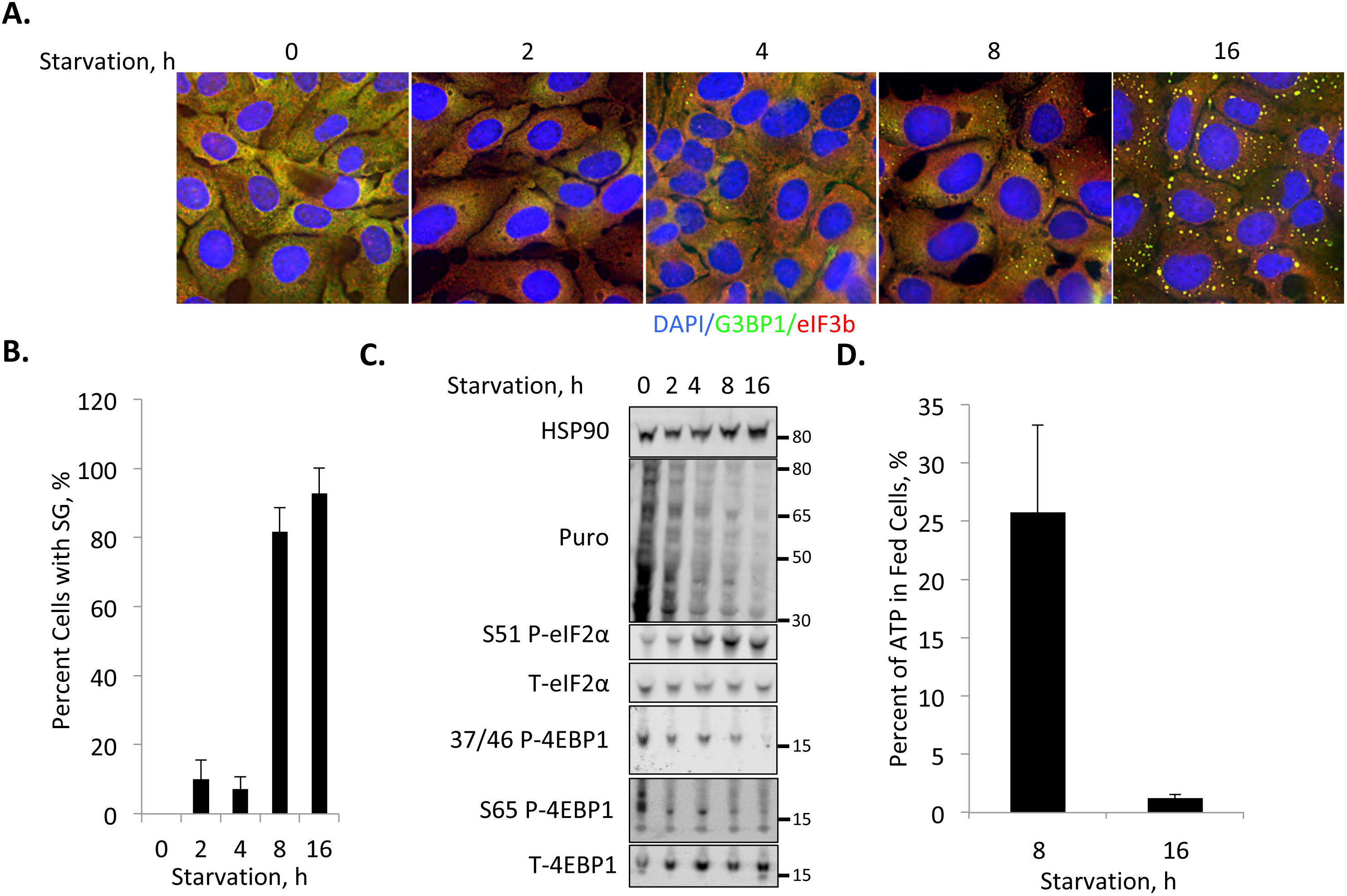
Chronic nutrient starvation induces stress granules with slow assembly kinetics. *A,* U2OS cells were starved of nutrients for the indicated times prior to fixation and staining with antibodies against G3BP1 (green) and eIF3b (red). *B,* Quantification of percent cells with SG from A. *C,* Western blot analysis of starvation stress kinetics. Puromycin labeling was used to analyze ongoing translation, and known pathways known to mediate translation repression are examined (mTORC1 and eIF2α), as indicated. Molecular weight standards are indicated. *D*, ATP levels during starvation at 8 and 16h time points. ATP is detected using a luciferase-based luminescence assay. ATP levels are expressed as a percent of control (fed) values at each time point. Results from all panels are representative of a minimum of three experimental replicates.

Although the kinetics of SG assembly during extreme nutrient starvation are slower than with acute stressors, the temporal relationship between eIF2α phosphorylation and SG assembly is reminiscent of a similar correlation during acute stress. This similarity suggests parallels may exist in these two contexts in regard to the molecular control of SG formation. To more thoroughly understand the composition of chronic SGs, we analyzed co-localization of other canonical SG proteins with G3BP1-containing SGs under nutrient deprivation. Similar to acute stress granules, we found co-localization of the RNA binding proteins HuR and Tia1 within nutrient starvation-induced SGs (Fig.2A). Also similar to acute stress-induced SGs, chronic starvation-induced SGs also stained positive for poly(A)^+^ RNA via fluorescence *in situ* hybridization (FISH) (Aulas et al. 2017). In contrast, the SGs induced by chronic starvation lack rps6 (Fig.2A) (Kedersha et al. 2002), suggesting that small ribosomal subunits are excluded from these structures. We therefore performed RNA FISH for 18S and 28S ribosomal RNA in cells subjected to both acute oxidative stress and chronic nutrient starvation (Fig.2B,C) (Kedersha et al. 2002; Kedersha et al. 2016; Sfakianos et al. 2018; Reineke et al. 2017; Tsai et al. 2016). Surprisingly, while 18S rRNA was strongly localized to arsenite-induced SG, 18S was weak or absent in nutrient starvation-induced SG (Fig.2B,C). We confirmed these results using a second 18S FISH probe, further examining 18s colocalization at an early time point (8 h) to determine if 18S signal concentrated within SG early when ATP levels were higher and then vacated at the late (16 h) time point (data not shown, Fig.2B,C). Consistent with previous work (Kedersha et al. 2002), 28S rRNA does not appear to concentrate in SGs in either context (Fig.2B).

**Figure 2:**
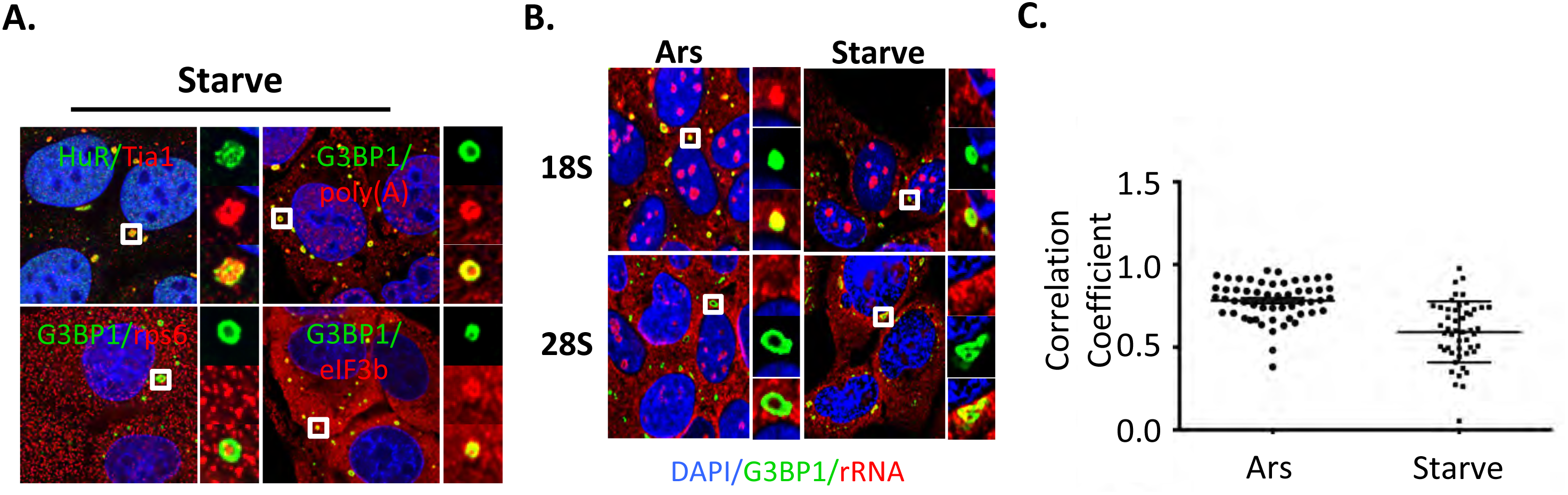
Starvation-induced SG have similar composition as other types of SG, but lack 40S ribosomes. *A,* U2OS cells were starved for 16h followed by fixation and staining for the indicated SG constituents. G3BP1 is the most well-studied RNA binding protein within SG; HuR and Tia1 are other well studied SG constituents; poly(A)^+^ mRNAs are included in canonical, acute stress granules; eIF3b is a translation initiation factor and its localization with SG defines SG; rps6 is a subunit of the 40S ribosome. *B,* RNA FISH was performed with probes directed against 18S and 28S rRNA after U2OS cells were exposed to either 250μM arsenite stress for 1h or chronic nutrient starvation (as indicated). *C,* Quantification of colocalization between 18S rRNA and G3BP1. Colocalization was measured using a Manders correlation test where 0 indicates no colocalization and 1 indicates perfect colocalization. Each data point represents the Manders coefficient for each cell spanning 8 fields per condition. Microscopy was performed with a 100X objective. A Student’s t-test was used to measure statistical significance (p<0.001, ^***^). Results from all panels are representative of a minimum of three experimental replicates.

Acute stress-induced SGs are highly dynamic and reflect a shifted equilibrium away from polysomes and towards SG formation. Trapping translation complexes in polysomes using pharmacological drugs such as cycloheximide and emetine shifts the equilibrium and induces disassembly of dynamic SGs (Kedersha et al. 2000; Aulas et al. 2017). The lack of colocalization of 18S rRNA with nutrient starvation-induced SG suggested these SGs are fundamentally different than previously described acute SGs, prompting us to test whether nutrient starvation-induced SGs would similarly disassemble in response to these drugs. As expected, arsenite-induced SGs were highly dynamic and could be disassembled when cells were co-treated with either cycloheximide or emetine (Fig.3A,B). Conversely, SGs assembling in response to nutrient starvation could not be disassembled with cycloheximide or emetine (Fig.3A,B).

**Figure 3:**
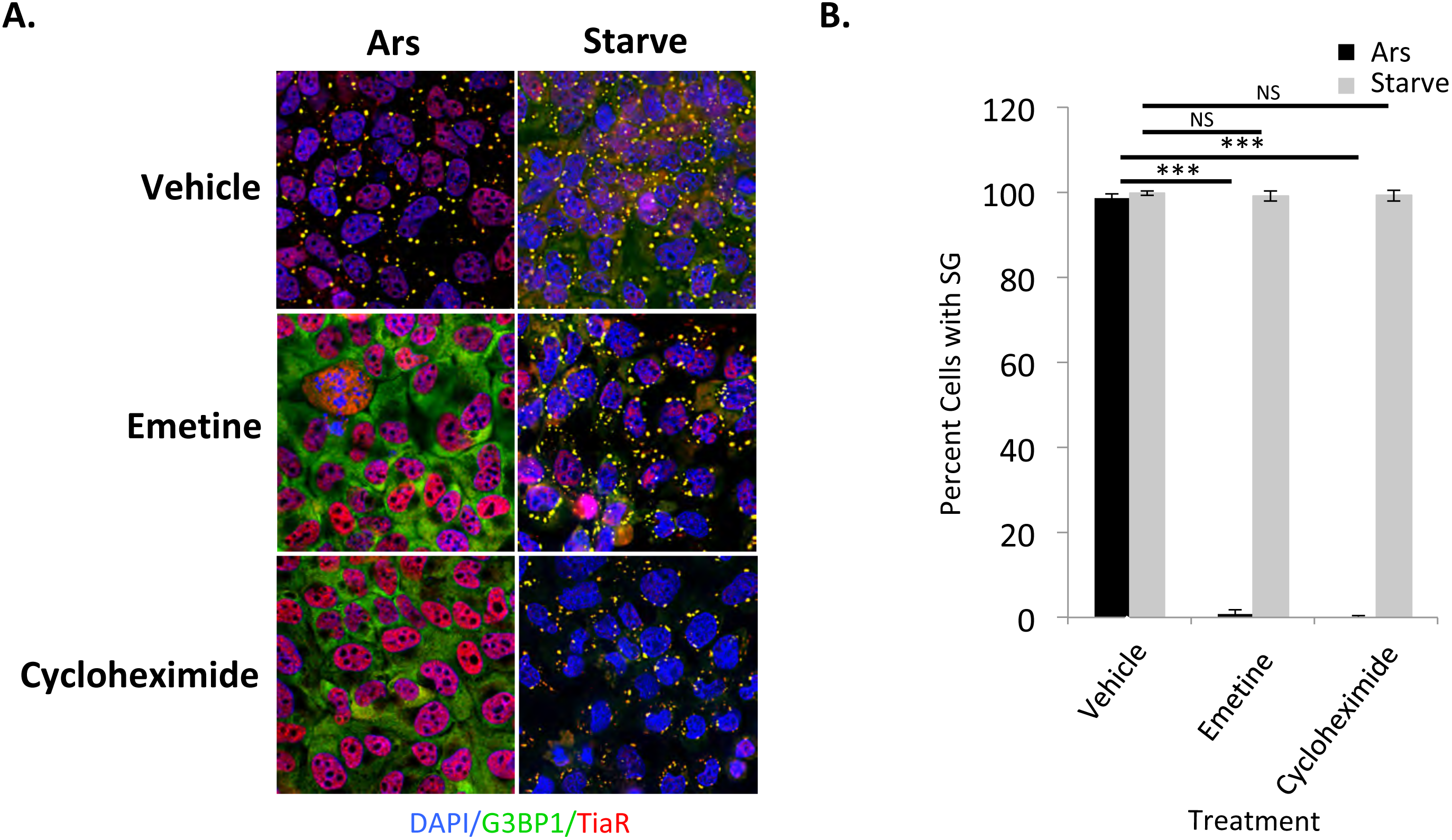
Chronic nutrient starvation induces static stress granules that do not exchange material with translating mRNPs. *A,* U2OS cells were either treated with 250μM arsenite for 1h or starved for 16h prior to fixation and staining with antibodies against G3BP1 (green) and TiaR (red). During the last hour of stress, 20μM emetine or 50μM cycloheximide were added to trap SG constituents within translating mRNPs by stalling elongating ribosomes on mRNAs to assess exchange of material with SG. *B,* Quantification of results shown in A. The y-axis depicts percent of cells with SG for each condition. Statistical analysis was performed with a Chi-test (NS indicates not significant and p<0.001,^***^). Results from all panels are representative of a minimum of three experimental replicates.

Because of the central role for eIF2α phosphorylation in SG induction for acute stressors, and the elevated eIF2α phosphorylation under these stress conditions (Fig.1C), we sought to formally test the importance of eIF2α phosphorylation in nutrient starvation-induced SG. To do this, we utilized mouse embryonic fibroblasts homozygous for an allele encoding an S51A mutant of eIF2α (AA/mutant), which cannot be phosphorylated (Guan et al. 2014; Reineke et al. 2012). Under nutrient starvation conditions, SG were not abundant in the AA MEFs, while control wild-type (SS/WT) were characterized by extensive SG formation (Fig.4A). While translation is still repressed in the AA MEFs during nutrient starvation, we were surprised to find that it is slightly elevated during stress as compared with the SS control MEFs (~35 % for AA vs 25 % for SS, Fig.4B). This result indicates that phosphorylation of eIF2α is critical for formation of SG precursor complexes, and other translation inhibition activities such as inhibition of eIF4f activity by mTORC1 are less important under these conditions. These results further indicate that one or more eIF2α kinases are likely to be activated under these stress conditions.

**Figure 4:**
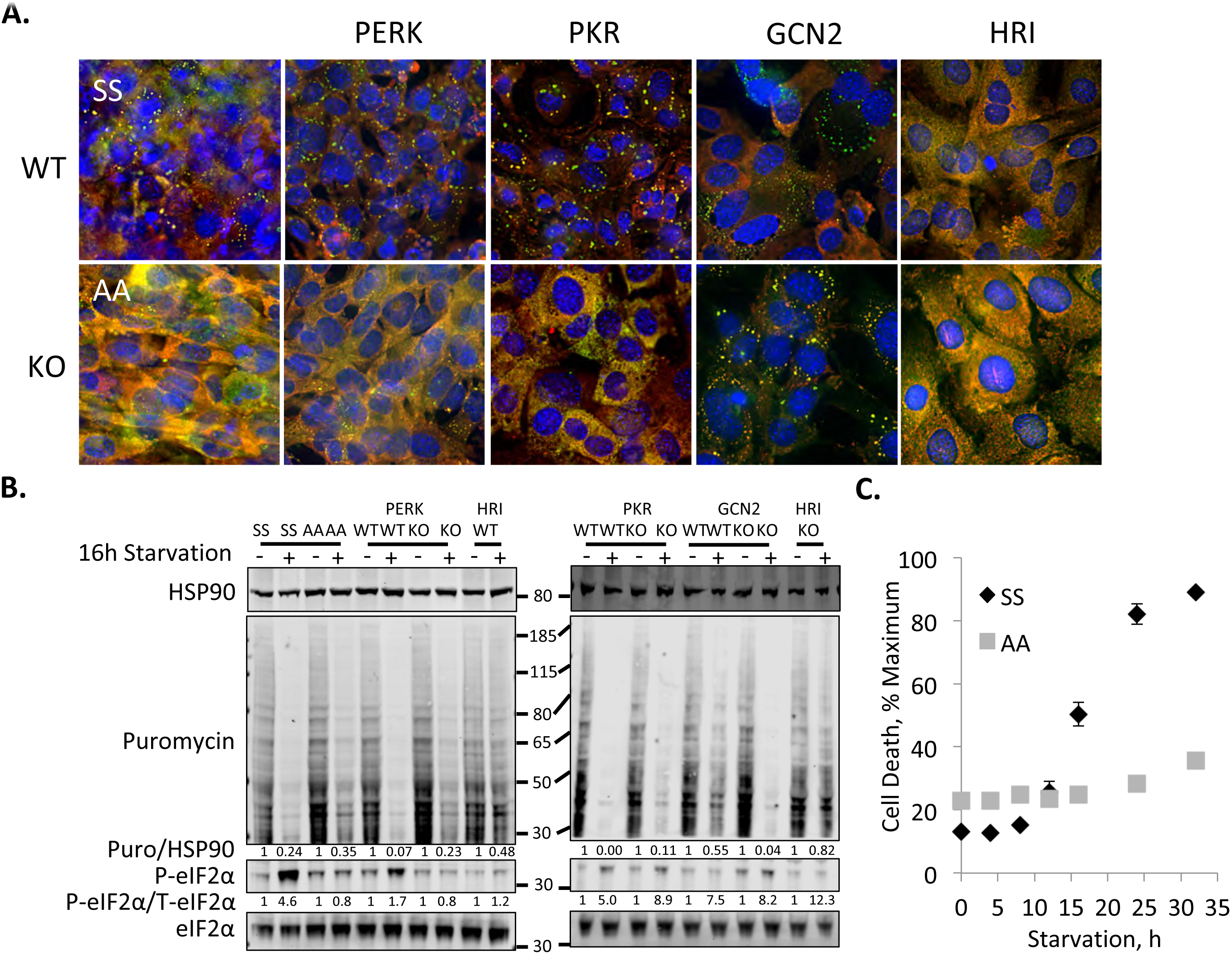
Role of eIF2α kinases in translation repression and SG induction during chronic nutrient starvation. *A,* The indicated eIF2α kinases-null or S51A eIF2α mutant mouse embryonic fibroblasts were stressed for 12 h prior to fixation and staining of SG with antibodies against G3BP1 (green) or eIF3b (red). *B,* MEFs were stressed as in A followed by Western blotting for the puromycin-labeled polypeptides to assess ongoing translation, or the indicated proteins. Molecular weight standards are indicated. Quantification of Western blots was performed using image studio for Li-Cor to avoid saturation effects, and ratios of total to phospho-proteins are indicated. *C,* Wild type and S51A eIF2α mutant MEFs were starved in the presence of ethidium homodimer-1 as described above. Ethidium fluorescence is normalized to 0 h and plotted against time. Results from all panels are representative of a minimum of three experimental replicates.

To determine whether specific eIF2α kinases are important in triggering SG assembly under chronic nutrient starvation, we utilized MEFs depleted of each of the previously described eIF2α kinases (HRI, PERK, PKR and GCN2). MEF genotypes were functionally validated using the distinct acute stressors that had been previously shown to selectively activate the various kinases (Fig.S1). In response to nutrient starvation, we monitored ongoing translation via puromycin pulse experiments, eIF2α phosphorylation levels, and SG assembly. Interestingly, PERK and PKR-null MEFs were characterized by reduced levels of SG formation (Fig.4A). These deficiencies both led to enhanced protein synthesis under the stress, but unlike PERK, PKR deficiency did not negatively affect eIF2α phosphorylation levels during nutrient starvation (Fig.4B), suggesting that PKR can act on SG components or assembly pathways independent of eIF2α phosphorylation. Depletion of GCN2 did not have a significant effect either on eIF2α phosphorylation or SG assembly (Fig.4A,B).

Using MEFs, we identified three factors important for SG condensation under chronic nutrient starvation: eIF2α phosphorylation, PERK and PKR. To test the contribution of these factors to cellular survival during chronic nutrient starvation, we performed a kinetic experiment in which SS and AA MEFs were subjected to nutrient starvation and cell death was monitored with ethidium homodimer staining of nuclei. The AA MEFs were remarkably less sensitive to cell death from nutrient starvation as compared to the wild-type SS counterparts (Fig.4C). We saw partial protection of both PERK and PKR-null MEFs during chronic nutrient starvation similar to our observations in the AA MEFs (Fig.S3).

These findings suggested the intriguing possibility that, in contrast to pro-survival acute stress-induced SG, some conditions induce pro-death SGs (Arimoto et al. 2008; Somasekharan et al. 2015). However, our experiments to this point had not yet excluded the contribution of ongoing translation and eIF2α phosphorylation as critical factors underlying the process of cell death here. Therefore, we attempted to test the importance of SG directly using U2OS cells in which the SG nucleating family members G3BP1 and G3BP2 had been inactivated using CRISPR/Cas9 (ΔΔ cells;(Kedersha et al. 2016)). We found that ΔΔ cells induce similar levels of eIF2α phosphorylation and translation repression during chronic nutrient starvation (Fig.5A). ΔΔ cells are incapable of SG formation under chronic nutrient starvation, indicating that G3BP1 and G3BP2 are key proteins for both acute and chronic stress granule formation (Fig.5B). However, we were surprised to find that the morphology of ΔΔ cell monolayers appeared considerably better under the chronic stress than the wild type U2OS control cells (Fig.5C). As such we evaluated cell death using annexin V/propidium iodide staining via flow cytometry, and found a significant reduction in dead and dying cells when the ΔΔ cells were compared with wild type (Fig.5D,E). When the same analysis was done on SS and AA MEFs, we saw that AA MEFs, which could not efficiently form SGs, had improved survival as well (Fig. 5F). Both U2OS and MEF cells appeared healthy and had low levels of dead cells under fed conditions (Fig.S4). Together these findings suggest SGs can function in a pro-death manner during chronic nutrient starvation, in contrast to previously described functions for SGs during acute stress.

**Figure 5:**
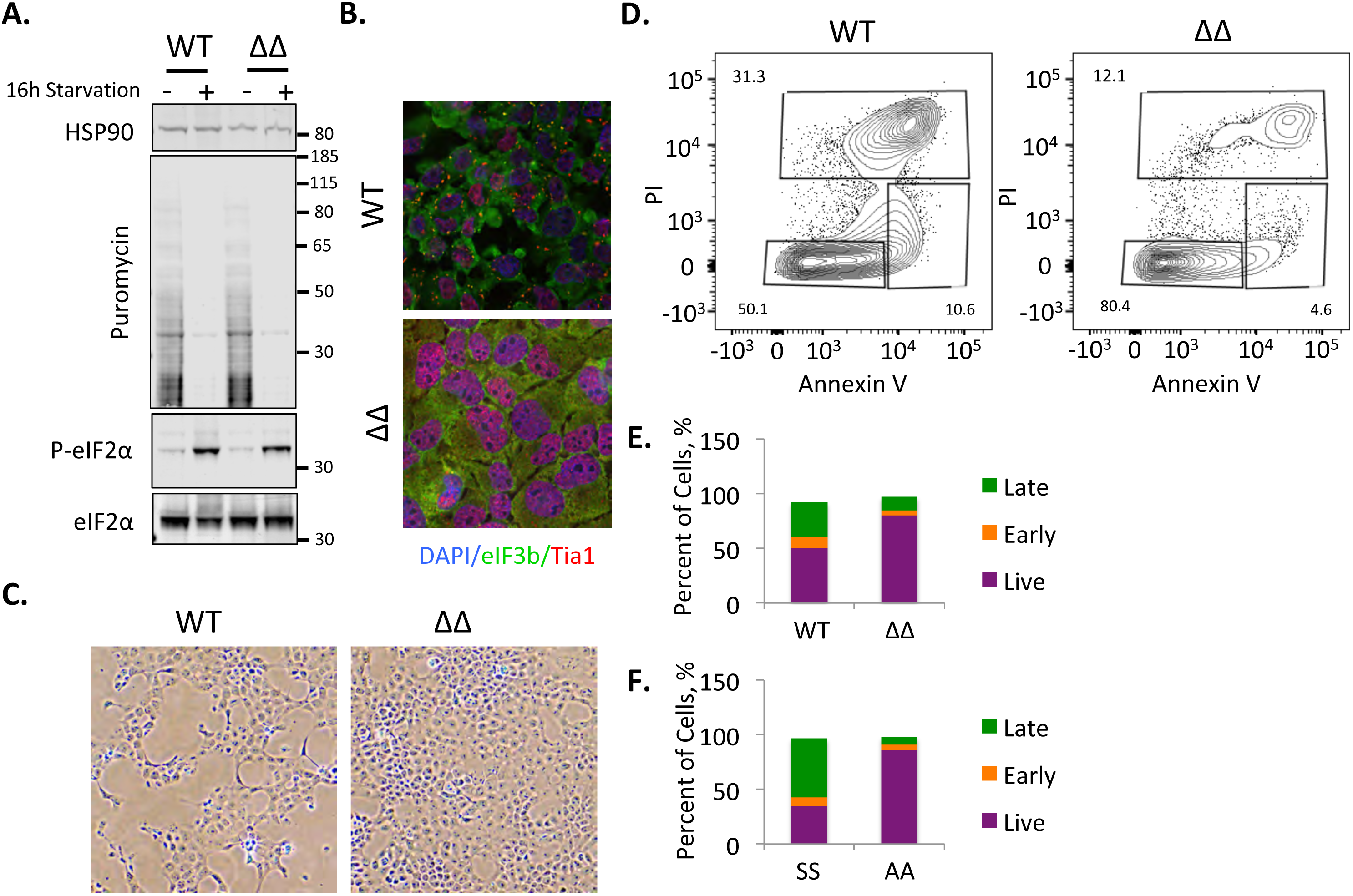
Chronic nutrient starvation induces pro-death SG. *A,* Wild type and G3BP1/2 double knockout U2OS cells were starved for 16h followed by western blotting for puromycin or the indicated proteins. Molecular weight standards are indicated. *B,* Cells treated as in A were stained for SG using eIF3b (green) and Tia1 (red) after 16h starvation. *C,* Phase contrast images taken with a 10X objective showing the appearance of the respective monolayers for wild type and ΔΔ U2OS cells. *D,* Flow cytometry was performed on fed and (see Fig.S4) starved U2OS cells stained with annexinV and propidium iodide to monitor the proportion of cells undergoing cell death. *E,* Graphical illustration of flow cytometry results on U2OS cells. The y-axis is percent of cells in each category with the total equaling 100%. *F,* Wild type and S51A eIF2α mutant mouse embryonic fibroblasts were subjected to the same analysis as in D and are represented as in F. Results from all panels are representative of a minimum of three experimental replicates.

## Discussion

We have characterized and delineated the importance of stress granules during chronic nutrient starvation for the first time. Nutrient starvation is a stress encountered by many organisms and cells in various disease states. While key pathways that regulate protein synthesis are well studied, their interplay under chronic stress conditions of multiple nutrients analogous to that experienced in disease is not understood. Under the nutrient starvation conditions described here, mTORC1 activity is dramatically downregulated; however, SG condensation is not visible until eIF2α is phosphorylated. Consistent with these kinetics, AA MEFs show strongly delayed SG formation despite robust translation repression. Protein synthesis is strongly repressed under all conditions examined here, although it is notable that protein synthesis is slightly elevated in stressed AA MEFs as compared to wild type SS control MEFs, as well as in PERK and PKR-null MEFs. These results indicate that while mTORC1 signaling kinetics do not appear to be important in SG assembly under these conditions, mTORC1 signaling is at least partly responsible for translation repression. Furthermore, the elevated level of protein synthesis in the AA MEFs during starvation suggests a subset of mRNAs continue to be translated in the presence of inhibited cap-dependent translation.

SG induction is dependent on eIF2α phosphorylation. This observation leads to the critical question of which eIF2α kinase(s) are activated under chronic nutrient starvation conditions. PERK activation appears to be very important for eIF2α phosphorylation under these conditions, resulting in impaired SG condensation. While PKR null MEFs still showed robust eIF2α phosphorylation during nutrient starvation, those cells did not robustly assemble SGs. This suggests PKR is important for SG condensation independent of its activity on eIF2α. We previously showed that SGs activate PKR in a pathway involving Caprin1, which could be important in maintaining phase separation of SG components under these conditions (Reineke et al. 2012; Reineke et al. 2015). These results indicate that PERK and PKR are important, through distinct pathways, for SG condensation during chronic starvation. PERK activation may suggest activation of an integrated stress response resulting from an unfolded protein response under these conditions, which will be the focus of future work. Interestingly, the integrated stress response downstream of the unfolded protein response is often associated with improved survival (Guan et al. 2017), but in this case it would suggest the integrated stress response downstream of PERK activation is associated with elevated cell death similar to systems in which sustained activation of the unfolded protein response is detrimental (Prola et al. 2017).

GCN2 is not activated under the nutrient starvation conditions used herein. This is not entirely surprising as the key amino acids that activate GCN2 are in sufficient abundance (Deng et al. 2002; Nikonorova et al. 2018). We cannot rule out HRI activation, as chronic nutrient starvation does induce oxidative stress, and oxidative stress was previously reported to induce SG through HRI activation (McEwen 2005). The wild type and HRI null MEFs were intrinsically resistant to SG under these stress conditions, which might be related to their slower growth characteristics as compared to the other MEF pairs. Even at late 24 and 32 h time points of nutrient starvation (Fig.S2), HRI wild type and knockout MEFs did not robustly display SG and were not dying in cell death assays. We do not believe HRI is a major regulator of eIF2α phosphorylation in the context of chronic nutrient starvation because the HRI-null MEFs had robust eIF2α phosphorylation (~12-fold, Fig.4B) under these conditions. However, strong HRI expression patterns are largely restricted to erythrocytes and macrophages (Chen 2014), suggesting any large role for HRI in these pathways would be highly context dependent.

We show that a novel type of SG is formed under starvation conditions depleted of glucose, glutamine, pyruvate and serum. These SGs are unique in that they do not contain 40S ribosomes, they are not dynamically exchanging with cytosolic components, and that they promote cell death. Stress granule formation has been suggested to be dependent upon ATP for assembly (Jain et al. 2016); however, nutrient starvation-induced SG assemble during times when ATP levels are low. Previous work indicated that inhibition of ATP production by addition of the non-hydrolyzable 2-deoxyglucose and CCCP inhibited assembly of acute oxidative stress granules (Jain et al. 2016). This difference may reflect the difference between acute and chronic stressors. We show that SG start appearing at 8 h of nutrient starvation when ATP levels are approximately 20% of fed control cells. By 16 hours when cells are beginning to die, ATP levels are a mere 1% of control cells. Therefore, the decreased dynamics of these SGs could be a result of impairing the machinery responsible for all or part of the movement of SG material. In the future we will investigate the importance of these aspects of translational control and SG function during chronic nutrient starvation. We will also attempt to understand the mechanisms of pro-death functionality of these SG.

Impaired cell death correlated with enhanced protein synthesis, although not at the levels of fed controls. We anticipated that enhanced translation during nutrient starvation would be detrimental to survival because cells are ATP depleted, and translation is a major consumer of cellular ATP (Stouthamer, 1973(Pontes et al. 2015)). Strikingly, we found that ongoing translation and the ability to prevent SG formation improved cellular survival during chronic nutrient starvation stress using the AA mutant MEFs (Fig.4C). We did not observe enhanced protein synthesis in ΔΔ U2OS cells during chronic nutrient starvation similar to findings during acute stress (Kedersha et al. 2016). This suggests that the observed elevation of protein synthesis during chronic stress is not a function of the absence of SGs. However, this possibility cannot be excluded and will be the subject of continued work.

## Methods

### Cell culture and treatments

All cells were cultured under standard conditions (5% CO_2_, 37 degrees Celsius) in DMEM with 10 % heat-inactivated FBS supplemented with 15 mM HEPES, pH 7.4). Nutrient starvation was performed in DMEM minus glucose, FBS, glutamine, and pyruvate for the indicated times. Nutrient starvation followed two washes in starvation medium. All cell lines used in this study were previously published (Kedersha et al. 2016; Reineke et al. 2012).

### Immunoblotting

Immunoblots were performed on lysates using NuPAGE MOPS gels (Thermo Fisher, Waltham, MA) after protein quantification with BCA assays (Thermo Fisher, Waltham, MA). Blocking was performed in TBST with 5% BSA. Immunoblots were conducted using the following antibodies: anti-HSP90 (BD Biosciences), anti-puromycin (Kerafast, Boston, MA, 3RH11), S51 P-eIF2α (Cell Signaling, Danvers, MA, 119A11), anti-eIF2α (Cell Signaling, Danvers, MA, D7D3), anti-T37/46 P-4EBP1 (Cell Signaling, Danvers, MA, 236B4), anti-S65 P-4EBP1 (Cell Signaling, Danvers, MA, 174A9), and anti-4EBP1 (Cell Signaling Danvers, MA, 53H11). Secondary antibodies are conjugated with Dylight dyes (Cell Signaling, Danvers, MA) and detected using the Li-Cor Odyssey imaging system (Lincoln, NE).

### Immunofluorescence Microscopy

Cells were seeded onto poly-D-Lysine-coated glass coverslips the day before starvation. Starvation proceeded in accordance with the above description followed by fixation in PBS with 4 % formaldehyde for 30 min. Cells were permeablized in PBS with 0.5 % triton x-100 followed by blocking with 5% BSA in PHEM buffer (Reineke et al. 2017). Cells were stained with antibodies against G3BP1 ((Reineke et al. 2012), or Santa Cruz, Dallas, TX, H10), eIF3b (Santa Cruz, Dallas, TX, N20), HuR (Santa Cruz, Dallas, TX, 3A2), Tia1 (Santa Cruz, Dallas, TX, C20), or rps6 (Cell Signaling, Danvers, MA, 5G10), and mounted with medium containing DAPI (Vector labs, Burlingame, CA). Imaging was performed as previously described (Reineke et al. 2017). Secondary antibodies were all raised in donkey and labeled with alexa488 or alexa 647 dyes (Thermo Fisher, Waltham, MA).

### *In situ* Hybridization

Cells were seeded as above and starved for the indicated time points. Cells were fixed in PBS with 4% formaldehyde for 15 min, immediately followed by fixation in ice cold methanol for 15 min. Cells were then processed as previously described (Aulas et al. 2017) using oligo(dT)_40_, 18S or 28S cy5-conjugated probes (Sigma-Aldrich, St. Louis, MO). rRNA probe sequences used were: 18S, TTGAGACAAGCATATGCTACTGGC; 28S, TAGGTTGAGATCGTTTCGGCCCCAAGACCTCTAATCATTCGCTTTACCGGATAAAACTGCGTGG. After RNA FISH, SG were stained using standard immunofluorescence procedures prior to mounting coverslips.

### ATP Analysis

ATP levels were detected with the luciferase-based Celltiter 2.0 reagent (Promega, Madison, WI) in accordance with the manufacturers protocol. Briefly, cells were seeded into 96-well plates in and either fed or starved for the indicated time points. The Celltiter reagent was mixed with cell culture medium at a ratio of 1:1, incubated 10 minutes and measured on a Tecan m200 plate reader (Mannedorf, Switzerland). After starvation, the cell monolayer in a parallel set of wells was stained with 0.5 % crystal violet as previously described (Feoktistova et al. 2016) to normalize luciferase values against. Absorbance of crystal violet was monitored at 590nm.

### Cell Death Analysis

Cell death was monitored on a Tecan m200 plate reader equipped for fluorescence measurements. Cells were seeded into a 96-well black-walled tissue culture-treated plate and grown overnight. Cells were then washed and starvation medium with 4 μM Ethidium homodimer-1 (Thermo Fisher, Waltham, MA) was added prior to measurement of fluorescence with 530/645 em/ex wavelengths. Measurements were then collected at the indicated time points. After 32 h, triton x-100 was added to each well to a final concentration of 0.5 % and incubated 15 min at 37^°^ C prior to reading ethidium homodimer-1 fluorescence to determine fluorescence when all cells are dead for normalization purposes.

### Flow Cytometry

Cells were starved as described above followed by harvesting the medium and trypsinization of the cells. Trypsinized cells were diluted in ice cold DMEM with 10 % FBS, and combined with starvation medium. Cells were then washed twice in PBS followed by resuspending cells at a concentration of 1 e^7^ ml^−1^ in annexin binding buffer containing 10 mM HEPES, pH 7.4, 140 mM NaCl and 25 mM CaCl_2_. Annexin V-FITC (BD biosciences, San Jose, CA) and propidium iodide (Roche, Indianapolis, IN) were then added and incubated 15min on ice prior to dilution of cells 5-fold in annexin binding buffer and filtration through mesh sieves. Single color and unstained controls were conducted for each experiment, with the single color annexin V-FITC control being treated 15min with 0.15 % bleach on ice prior to staining. Flow cytometry was performed on an LSRII instrument (BD biosciences, San Jose, CA) at the Cytometry and Cell Sorting Core Facility at Baylor College of Medicine.

### Colocalization and SG Quantification, and Statistics

Colocalization analysis was performed using a custom-made Matlab script to measure the Manders Colocalization coefficients (MCC) (Manders *et al,* 1993). Briefly, nuclei were segmented in the DAPI channel using a watershed algorithm. Cell boundaries were then approximated by dilating the nuclear mask. Within each cell, G3BP1 (SG) and 18S rRNA signal were segmented by intensity thresholding using the Otsu method. MCCs between each channel (G3BP1 (green, G), and 18S rRNA (red, R)) were then calculated as follows:

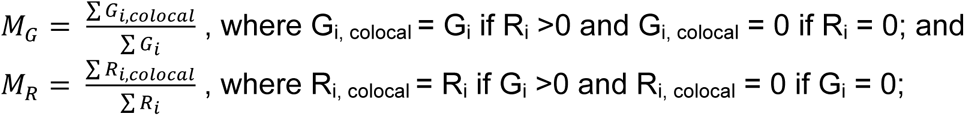

Only the M_G_ is shown for simplicity, and because the 18S signal was diffuse resulting in only a small fraction colocalizing with SG even under arsenite-treated conditions.

For quantification of percent of cells with SG, Image J was used. For this analysis, cell nuclei was counted after thresh holding with the analyze particles function. Subsequently, G3BP1 signal was detected with the detect local maxima function as previously described (Reineke et al. 2017). This was done in at least four fields per condition representing over 200 cells. To determine the percent of cells with SG, cells devoid of SG were manually counted in each field and tallied against the total number of cells. A Chi-test was performed for percent cells with SG, and a Student’s t-test was performed on Mander’s correlation data. Each experiment was repeated at least 3 times.

## Acknowledgements

The authors would like to thank Drs. Nancy Kedersha and Maria Hatzoglou for generous contributions of cell lines. This work was supported by an American Cancer Society - Athena Water Breast Cancer Research Scholar Grant (RSG-15-088-01RMC) to J.R.N., and NCI CA190467 to J.R.N. This project was supported by the Cytometry and Cell Sorting Core at Baylor College of Medicine with funding from the NIH (CA125123 and RR024574) and the expert assistance of Joel M. Sederstrom, as well as the Integrated Microscopy Core with funding from P30 Cancer Center Support Grant (NCI-CA125123), P30 Digestive Disease Center (NIDDK-56338-13/15), CPRIT (RP150578), John S. Dunn Gulf Coast Consortium for Chemical Genomics.

## Supplementary Figures

**Figure S1:**
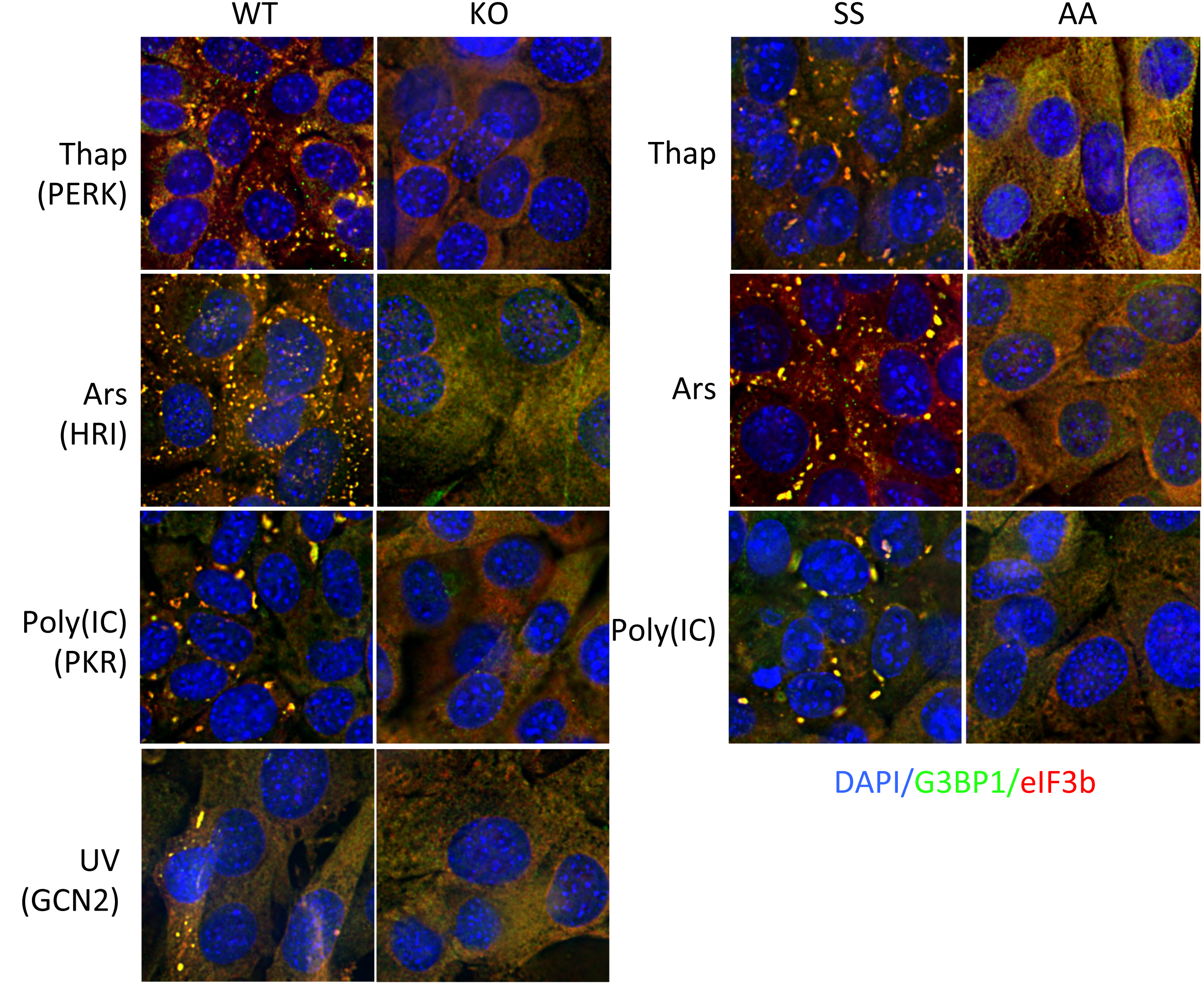
EIF2AK wild type and knockout MEFs were characterized to confirm they respond in accordance with known properties reflecting genotype. Each EIF2AK wild type and KO MEF pair was exposed to the indicated stress (2μM thapsigargin for PERK and S51A MEFs, 2h; 250μM arsenite, 1h for HRI and S51A MEFs; transfection of 100ng poly(IC), 6h for PKR and S51A MEFs; or 200J/m2 UV, 2h recovery for GCN2 MEFs) followed by staining for G3BP1 (green) and eIF3b (red) to mark SG. Several fields of cells per condition were acquired, and representative images are shown.

**Figure S2:**
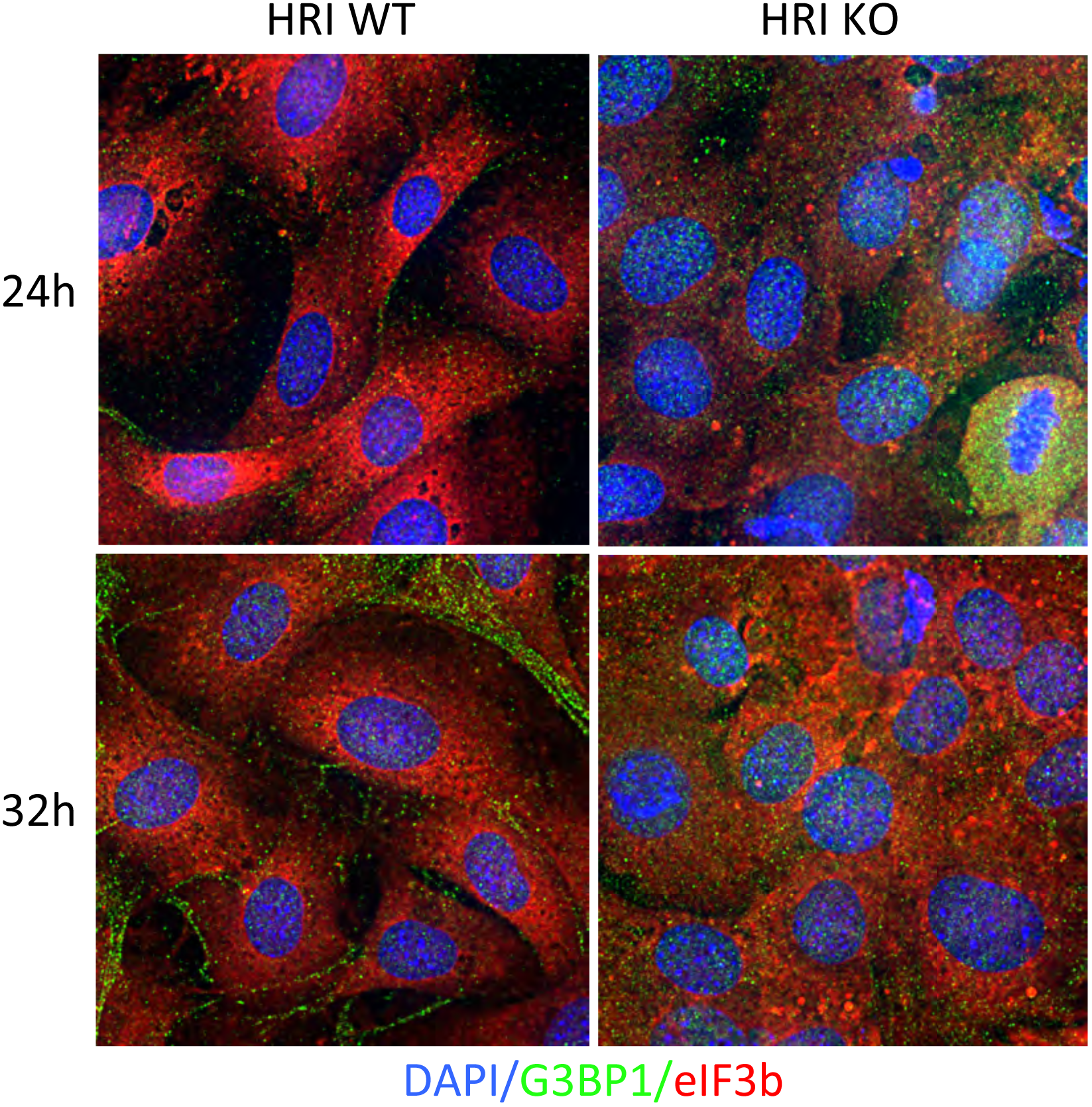
HRI wild type and knockout MEFs are intrinsically resistant to SG induction. HRI wt and KO MEFs were stained for the SG markers G3BP1 (green) and eIF3b (red) at 24 and 32h of starvation. No SG are visible. Red puncta represent some type of puncta that are extracellular as indicated by staining outside of the cell area across multiple fields.

**Figure S3:**
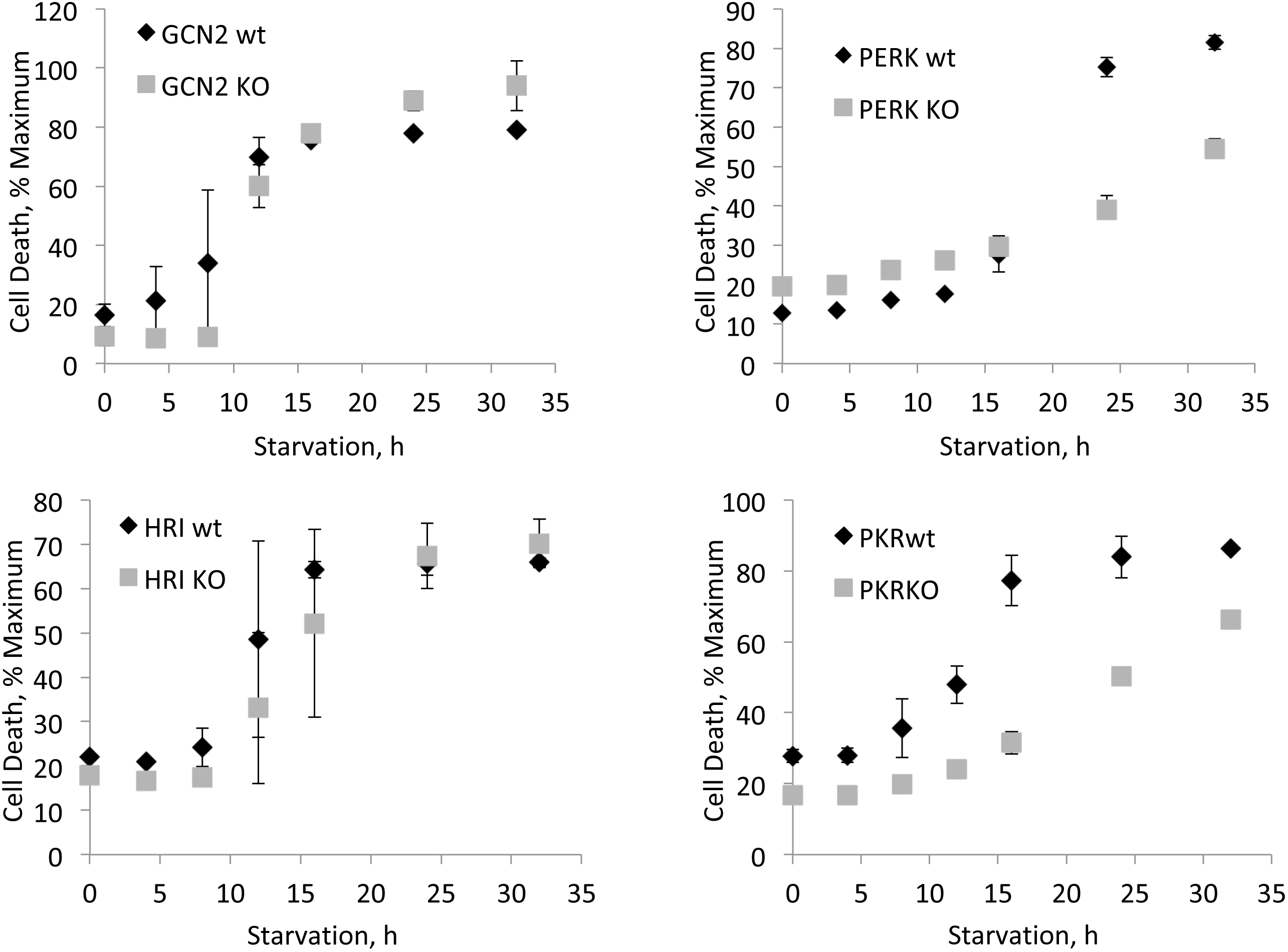
Survival Curves of EIF2AK wild type and knockout MEFs. Cells were starved for the indicated time points in the presence of ethidium homodimer-1. Time points were measured with a fluorescent plate reader as described in the Methods. Black diamonds represent wild type and gray squares represent paired KO cell lines.

**Figure S4:**
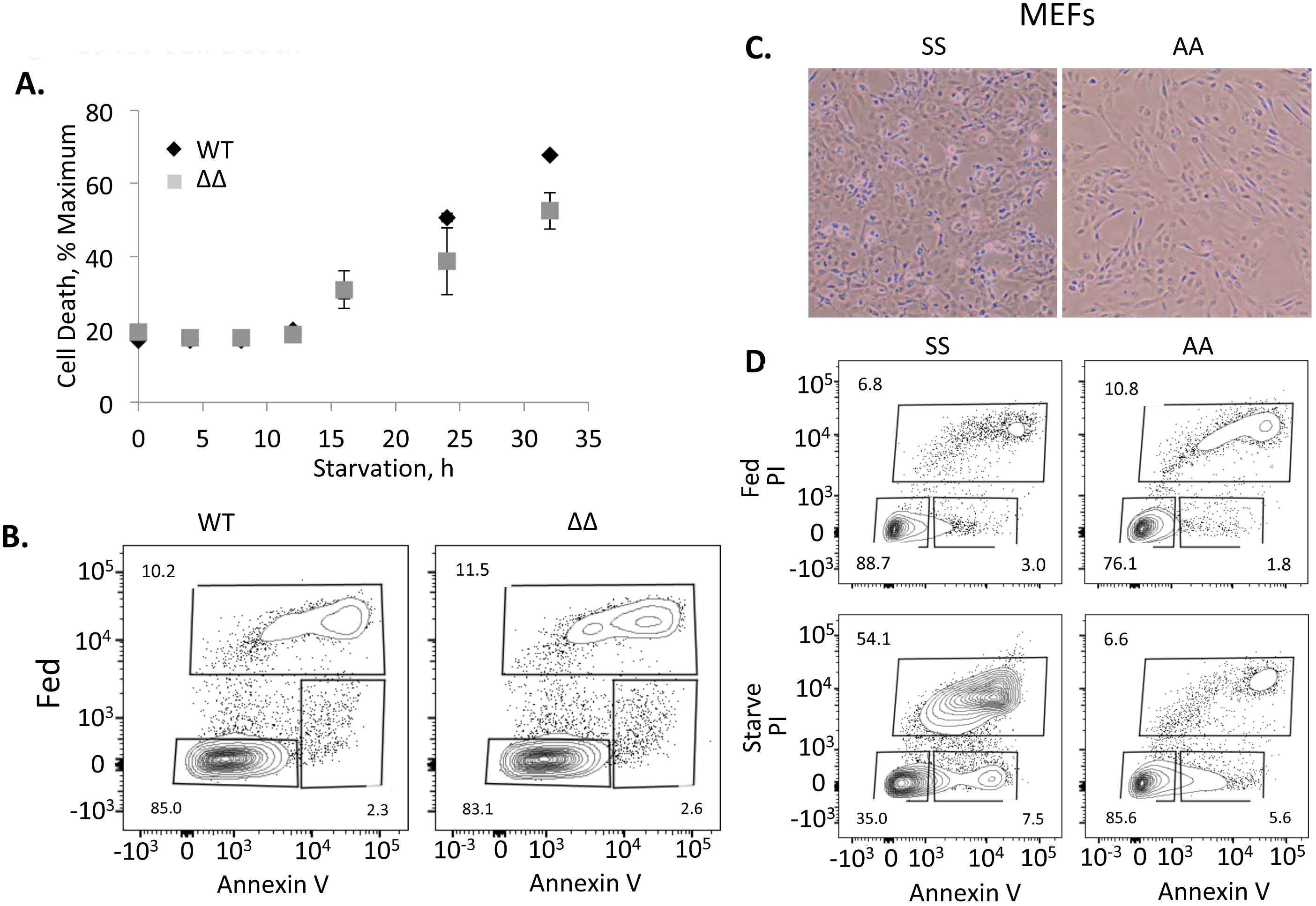
Cell death analysis to compliment Figure 5. *A,* Ethidium homodimer-1 staining of wt and ΔΔ U2OS cells as described in Figure S2 to show similar results as those shown by cell monolayers and flow cytometry. *B,* Flow cytometry with AnnexinV and propidium iodide staining of fed wt and ΔΔ U2OS showing that cells are healthy. *C,* Cell monolayer for SS and S51A MEFs showing that more dead/dying cells are present in SS MEFs as compared to the S51A mutant consistent with ethidium homodimer-1 staining and flow cytometry. *D,* Flow cytometry with AnnexinV and propidium iodide staining of wild type (SS) and S51A (AA) MEFs. This data was repeated thrice and is summarized in Figure 5f.

